# PRDM9 and an Epidemic of Gene Conversion and Non-Homologous Recombination among *Alu* Elements in Ancestral Gorillas

**DOI:** 10.1101/241356

**Authors:** Aaron C. Wacholder, David D. Pollock

**Author notes:** Corresponding Author*: David D. Pollock, Department of Biochemistry and Molecular Genetics, University of Colorado School of Medicine, Aurora, CO, 80045 USA. *Email:.

## Abstract

We performed a genome-wide scan for recombination-mediated interlocus gene conversion and deletion events among a set of orthologous *Alu* loci in the Great Apes, and were surprised to discover an extreme excess of such events in the gorilla lineage versus other lineages. Gorilla events, but not events in other Great Apes, are strongly associated with a 15 bp motif commonly found in *Alu* sequences. This result is consistent with evolutionarily transient targeting of the motif by PRDM9, which induces double strand breaks and crossovers during meiosis at specific but rapidly changing sequence motifs. The motif is preferentially found in conversion recipients but not donors, and is substantially depleted in gorillas, consistent with loss of PRDM9 targets by meiotic drive. Recombination probability falls of exponentially with distance between loci, is reduced slightly by sequence divergence, and drops substantially with recipient divergence from the target motif. We identified 16 other high-copy motifs in human, often associated with transposable elements, with lineage-specific depletion and nearby gene conversion signatures, consistent with transient roles as PRDM9 targets. This work strengthens our understanding of recombination-mediated events in evolution and highlights the potential for interactions between PRDM9 and repetitive sequences to cause rapid change in the genome.

## Main

During meiosis, double strand breaks are induced in chromosomes to initialize recombination.^1^ In mammals, specific sites in the genome are targeted for these breaks by binding of the protein PRDM9, resulting in recombination hotspots at these sites.^1,2^ Though recombination typically occurs between allelic pairs at the same locus, it can also occur ectopically between homologous sequences at distinct loci. Such interlocus recombination has major consequences for genome evolution, as crossovers between homologues at different loci can lead to segmental deletions and duplications. Furthermore, it can also result in interlocus gene-conversion, a process that involves non-reciprocal transfer of genetic information from one homologue to another. Allelic gene conversion contributes to substitution rate variation and GC-content heterogeneity in a variety of taxa^3,4^, and interlocus gene conversion drives concerted evolution among gene families^5^.

To better understand the dynamics of interlocus recombination-mediated events (RMEs) in primates, we performed a genome-wide scan for RMEs among *Alu* elements in the Great Apes. We searched for two possible consequences of such events, interlocus gene conversion and *Alu* recombination-mediated deletion (ARMD). To identify and characterize gene conversion events, we developed a novel method, TEConv, that uses a Bayesian phylogenetic approach. We applied this method to *Alu* loci orthologous among four Great Apes, identifying gene conversion events on each branch of the phylogenetic tree leading to these four species. For each conversion event, TEConv infers the lineage on which the event occurred, identifies both the donor and recipient loci, and probabilistically infers the sequence at each locus before and after the conversion. Conversion was tested between pairs of elements within 100 kb of each other, and false positive rates were estimated by placing each element at a random position in the genome at least 1 Mb from its true position (where there are likely to be essentially no true conversion events) and re-running the algorithm. Thresholds for inclusion into a high-confidence gene conversion set were specified to obtain a 5% false positive rate. Batzer et al. previously identified ARMD events in the human^6^ and chimpanzee^7^ lineages; here, we replicated this work using similar methodology and extended it to the gorilla and orangutan.

Surprisingly, the gorilla lineage shows a large excess of both gene conversion and ARMD relative to the other Great Apes (Table 1). Of 2,551 gene conversion events identified, nearly all (2514) occurred along the gorilla lineage, and over 4 times as many ARMD were found in the gorilla as in any other lineage considered. It appears that the terminal gorilla branch experienced vastly higher rates of RMEs among *Alu* elements than human, chimpanzee or orangutan. This could be explained if PRDM9 evolved to bind *Alu* elements in gorilla ancestors, but not those of other Great Apes. Because sequences that bind strongly to PRDM9 are preferentially converted by sequences that bind less strongly, PRDM9-binding motifs deplete rapidly from the genome, driving the extremely rapid evolution of PRDM9 to find new targets^8–10^. As a result, there is essentially no overlap in recombination hotspots between human and chimpanzee, so lineage-specific PRDM9 binding is not surprising^11^. We reasoned that if a transient PRDM9 mechanism explains excess RMEs associated with gorilla Alus, there might be a common *Alu* sequence motif that is associated with RMEs in the gorilla lineage. However, currently-active PRDM9 motifs have been identified by analyzing recombination hotspots^2^, and *Alu* gene conversions in the gorilla lineage are not particularly close to modern gorilla hotspots^12^: 12.4% of gene-converted *Alu* loci are within 10 kb of a hotspot (318 out of 2,571), compared to 13.5% of non-converted loci (34,021 out of 251,496). This suggests that any PRDM9 motif associated with excess gorilla *Alu* recombination is different from current motifs and no longer active.

**Table 1.**
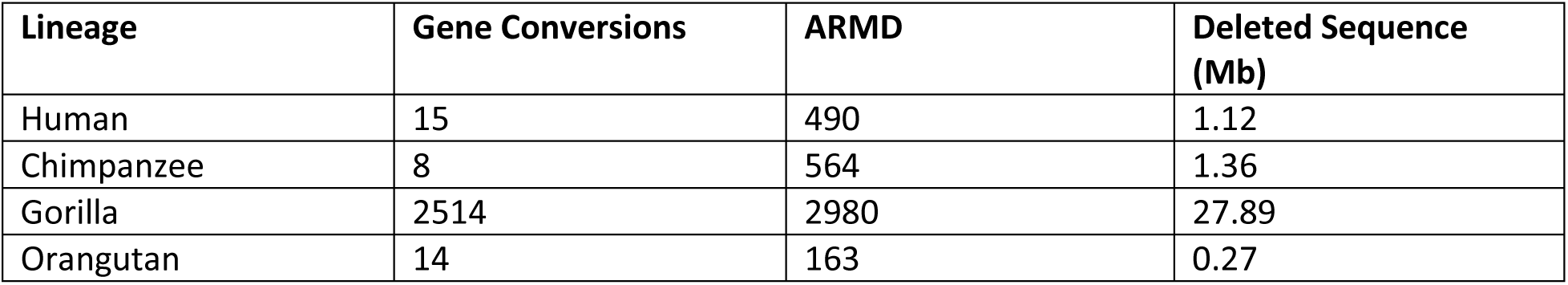

To search for this motif, we analyzed frequencies of pre-conversion variants at each site in our set of *Alu* sequences, comparing elements that underwent gene conversion in gorilla with elements that did not. The inferred state of each sequence in the common ancestor of Great Apes was chosen as the best available indicator of the pre-conversion sequence. Among two major classes of *Alu, AluY* and *AluS*, there is a region from positions 242–256 where the most common nucleotide variants are substantially elevated in conversion acceptors compared to unconverted elements (Figure 1). Even a single difference from the most common motif in this region is sufficient to decrease the relative probability of conversion at a locus by about 3-fold (Figure 2); additional differences reduce the probability further (Figure 3). In contrast, the state of this putative binding motif does not strongly impact the probability that a sequence will act as a donor element (Supplementary Figure 1). For ARMD events, we cannot distinguish the roles of the two recombination participants, but we can compare the sequences of pre-ARMD elements (inferred at the root of the Great Apes) to other elements. Gorilla *Alu*S elements with an intact motif were 1.38 times more likely to have been involved in ARMD events than other gorilla elements (95% credible interval: 1.30–1.57), while human and chimpanzee *Alu*S elements with the motif do not show evidence of increased recombination probabilities (95% credible intervals: 0.89–1.22 and 0.93–1.30, respectively), as expected if it were a transient PRDM9 binding motif.

**Figure 1.**
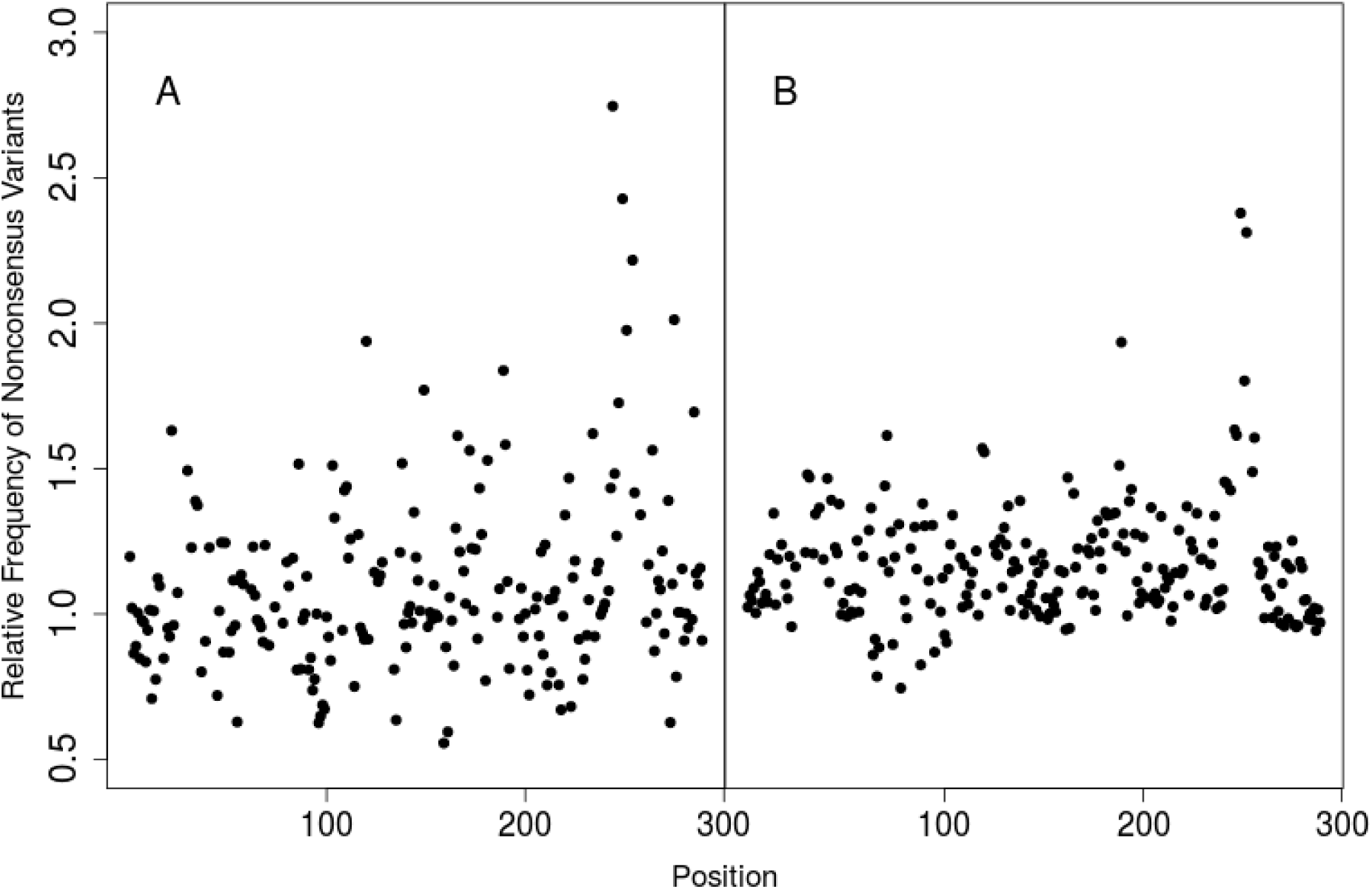
Ratio of the Frequencies of Uncommon Variants for Unconverted Elements Versus those for Gene Conversion Recipient Elements, for each Position in *Alu*. The proportion of *Alu* loci with uncommon variants at the root of the Great Apes was calculated for each site, and the ratio of counts for *Alu* elements that were not converted on the gorilla lineage versus those that were conversion recipients is plotted for each site. Positions differing between *Alu*S, *Alu*Y, and *Alu*J were excluded. Analysis was conducted separately for A) *Alu*Y and B) *Alu*S elements.

**Figure 2.**
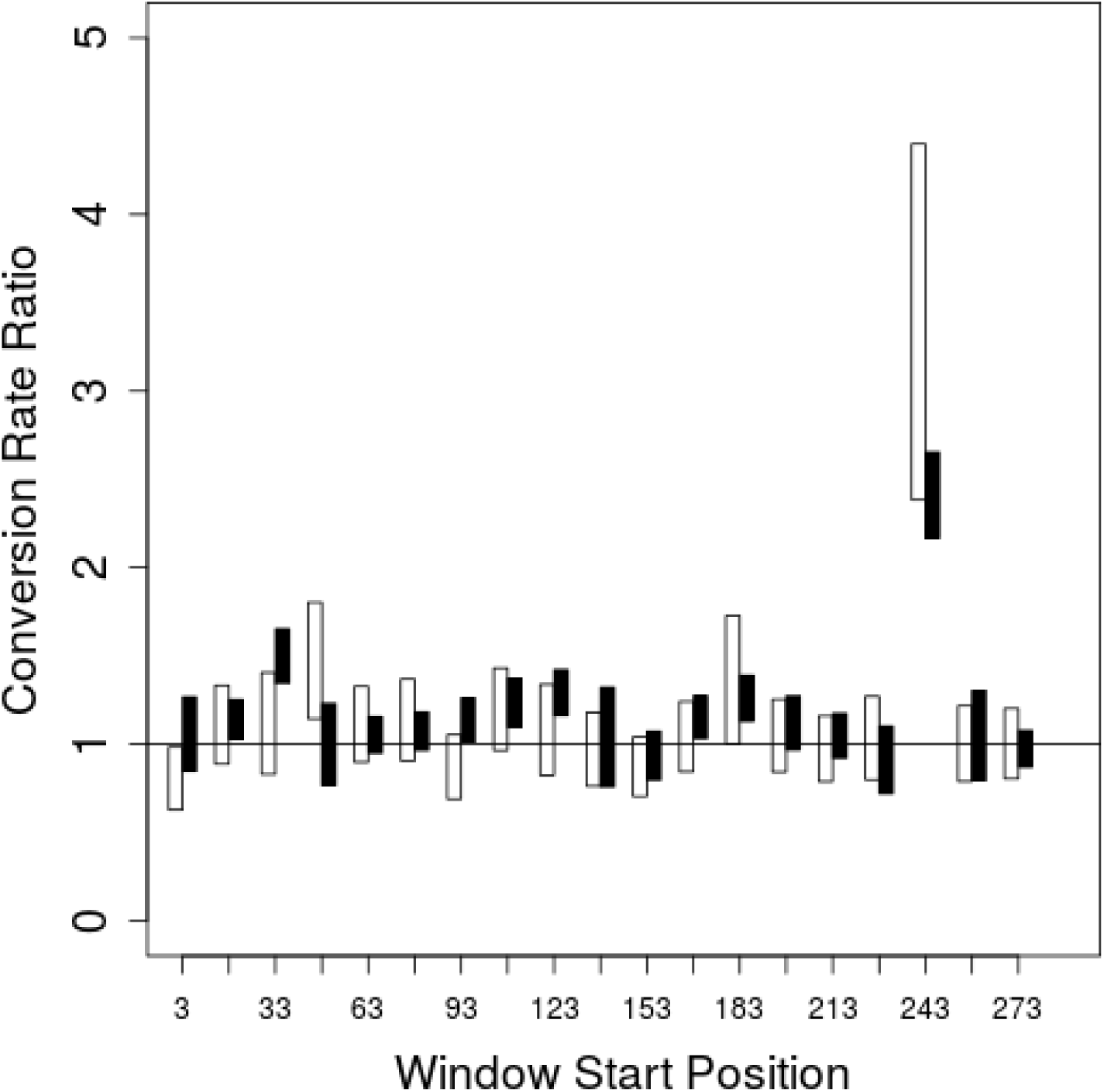
Conversion Acceptor Ratio Between Elements with and Without Perfect Match to the Most Common Sequnce in 15 bp Windows Across *Alu*. For 15 bp windows across *Alu*, the proportion of elements that were determined to be conversion recipients on the gorilla lineage was calculated for loci that had perfect match to the consensus at the root of the Great Apes in that window (excluding positions differing between *Alu*S, *Alu*Y, and *Alu*J) versus all other elements. The white bars show 95% credible regions for *Alu*Y elements, while the black bars show 95% credible regions for *Alu*S elements.

**Figure 3.**
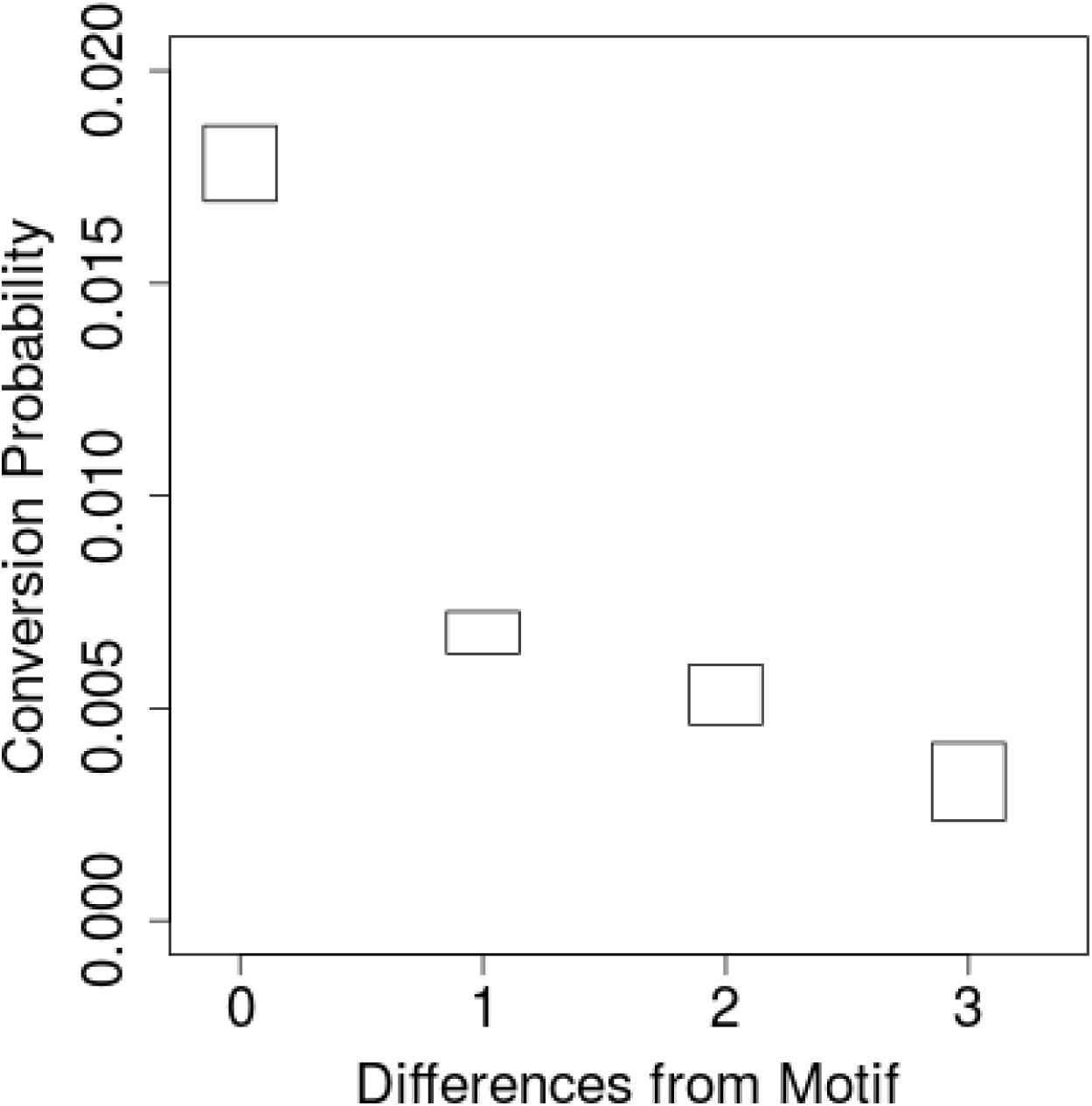
Conversion Probability by Motif Differences. The conversion recipient probability was estimated dependent on number of differences from the putative PRDM9 binding motif. Boxes show 95% credible intervals.

As expected of a PRDM9 binding motif, the putative motif we identified is depleted among *Alu* elements in gorillas relative to other Great Apes, while motifs differing by 1 or 2 nucleotides from the ancestral motif are more frequent (Supplementary Figure 2). The gorilla genome has 62,138 *Alu* copies that match the motif while other Great Apes have between 68,245 and 69,456 copies with intact motifs. Identified interlocus gene conversion events explain 11% of the approximately 6,000 motif copies lost in gorilla compared to other great apes; the remaining losses can be explained by unidentified interlocus conversion events or biased allelic gene conversion.

The distance between the midpoints of each pair of *Alu* loci involved in gene conversion events in gorilla, and in ARMD events in all four Great Apes, was measured (there were two few conversion events in the other Great Apes for analysis). The rate of interlocus recombination appears to decline exponentially with distance between the loci (Figure 4). Gene conversion and ARMD events in the gorilla lineage show similar distance distributions to each other, as expected based on a common PRDM9-binding double-strand break mechanism. In contrast, average gorilla ARMD distances were much greater than in other Great Apes, suggesting the possibility of a different mechanism for *Alu*-mediated interlocus recombination when *Alus* are not specific PRDM9 binding targets. As a consequence of larger average deletion size, while gorilla experienced only around five times as many ARMD events as human or chimpanzee, it lost over 30-fold as much sequence as either of those lineages (Table 1).

**Figure 4.**
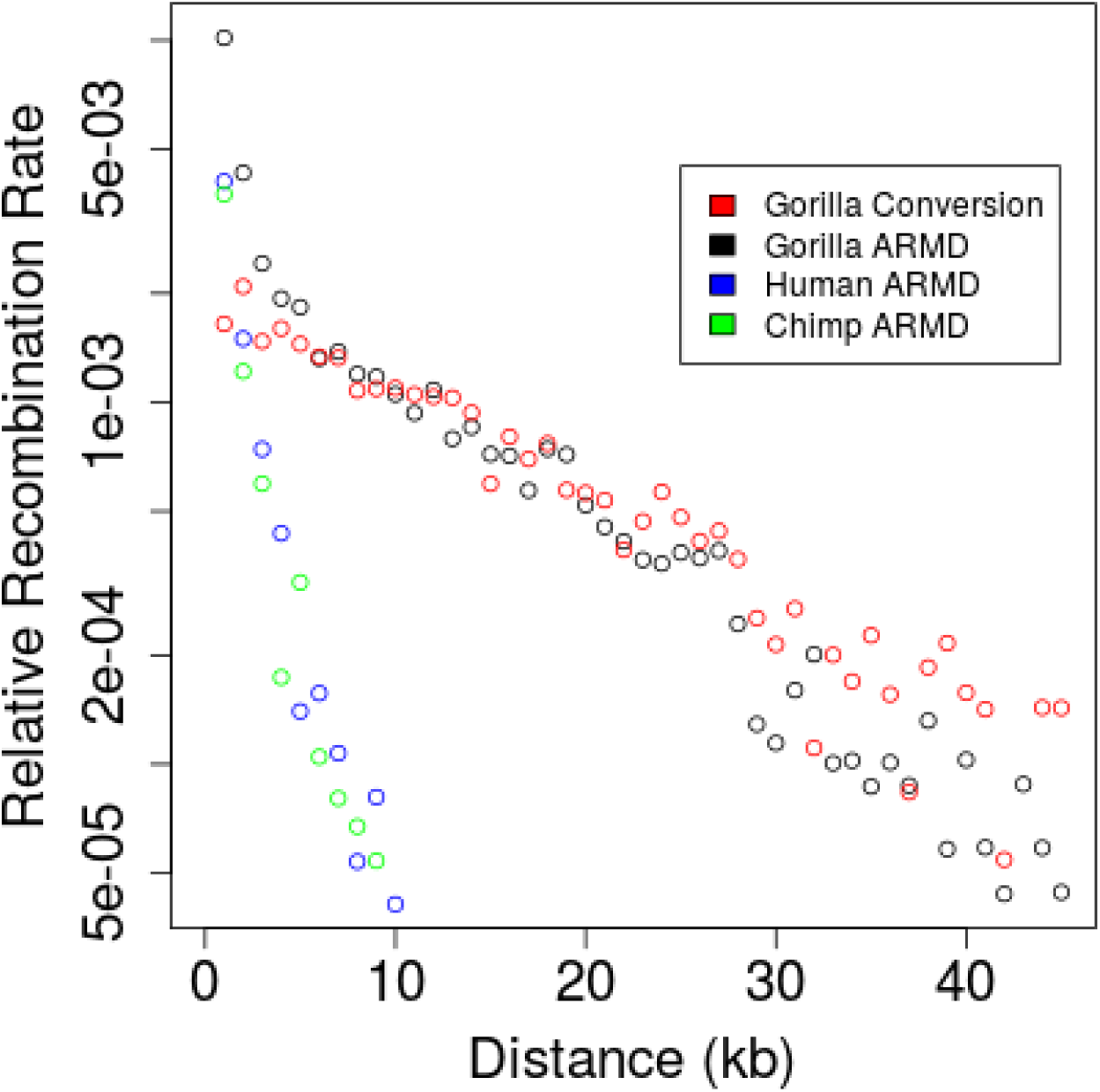
Recombination Rate by Distance. All possible *Alu* pairs were placed in bins of 1,000 bp based on the distance between inferred gorilla lineage conversion donors and recipients, with distance measured at the *Alu* midpoint. The proportion of possible pairs in each bin that were in the gorilla gene conversion set or were identified as ARMD participants on the human, chimpanzee, or gorilla lineage is plotted.

The posterior estimated mean interlocus conversion tract size was 118 bp (median 109 bp, standard deviation 54 bp). Tract size frequency among identified conversion events rises rapidly from 25-75 bp, then slowly declines in the range from 100-200 bp (Figure 5). To correct for bias against detecting shorter tracts, conversion events were simulated and the false negative rate for each tract size was used to infer the corrected distribution including missing events; this does not greatly change the general pattern (Figure 5). We also considered the number of times each position in the *Alu* element was covered by a conversion tract (Figure 6). The most covered region of the element is in the middle of the 289 bp *Alu* alignment, around positions 136-166, and the most covered site, position 149, is included in 65% of tracts. In contrast, the ends are covered much less: for example, position 20 is in only 5% of tracts, and position 269 in 17% of tracts. The putative PDRM9 motif at *Alu* alignment positions 242-256 does not appear as an outlier in conversion tract coverage (only 36% of tracts cover the motif).

**Figure 5.**
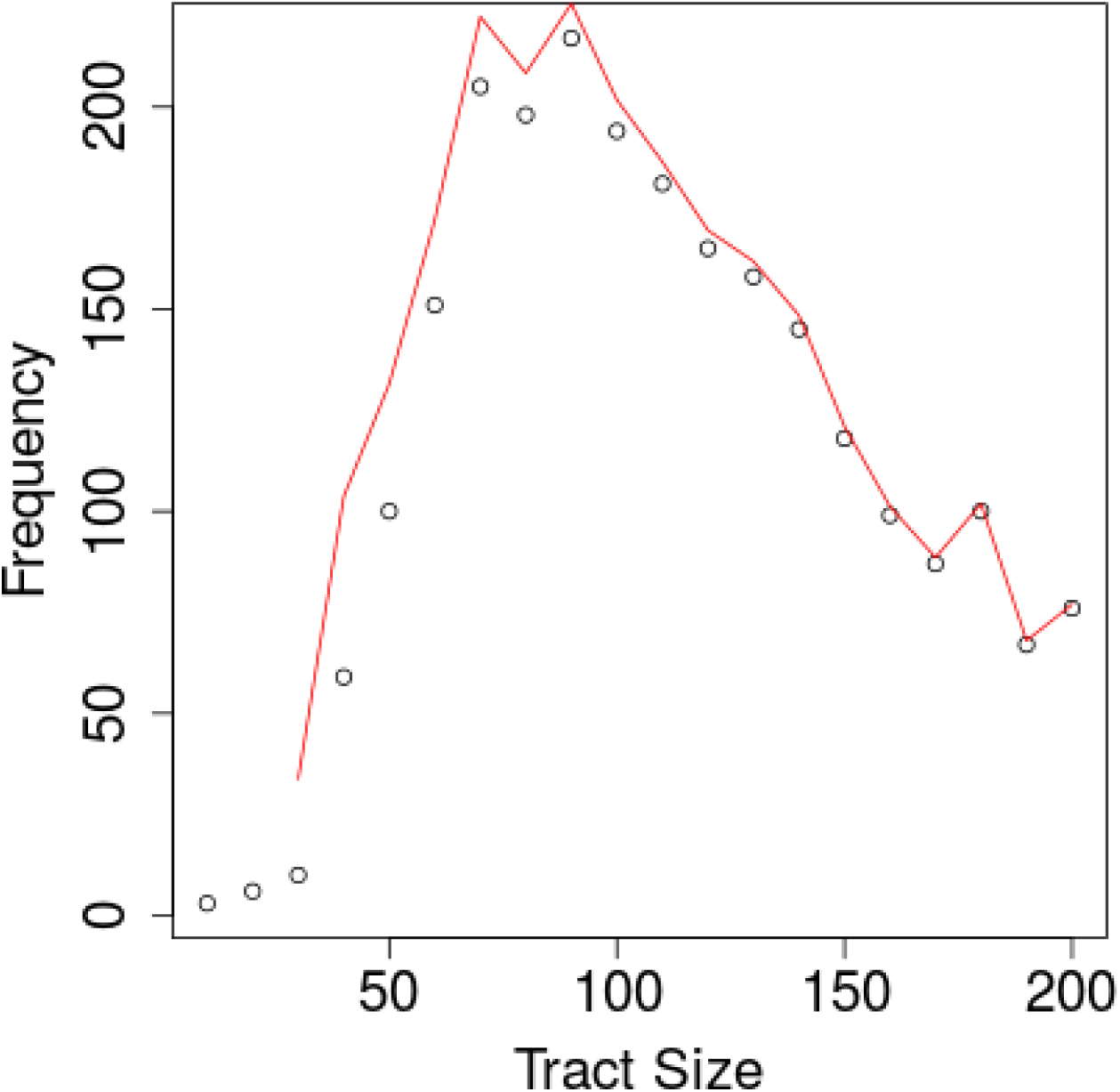
Tract Size Frequencies. All pairs in the conversion set were placed in 10 bp bins based on tract size, and the number of pairs in each bin was plotted. The red line gives the estimated frequency in each bin after adding in inferred missed conversion events.

**Figure 6.**
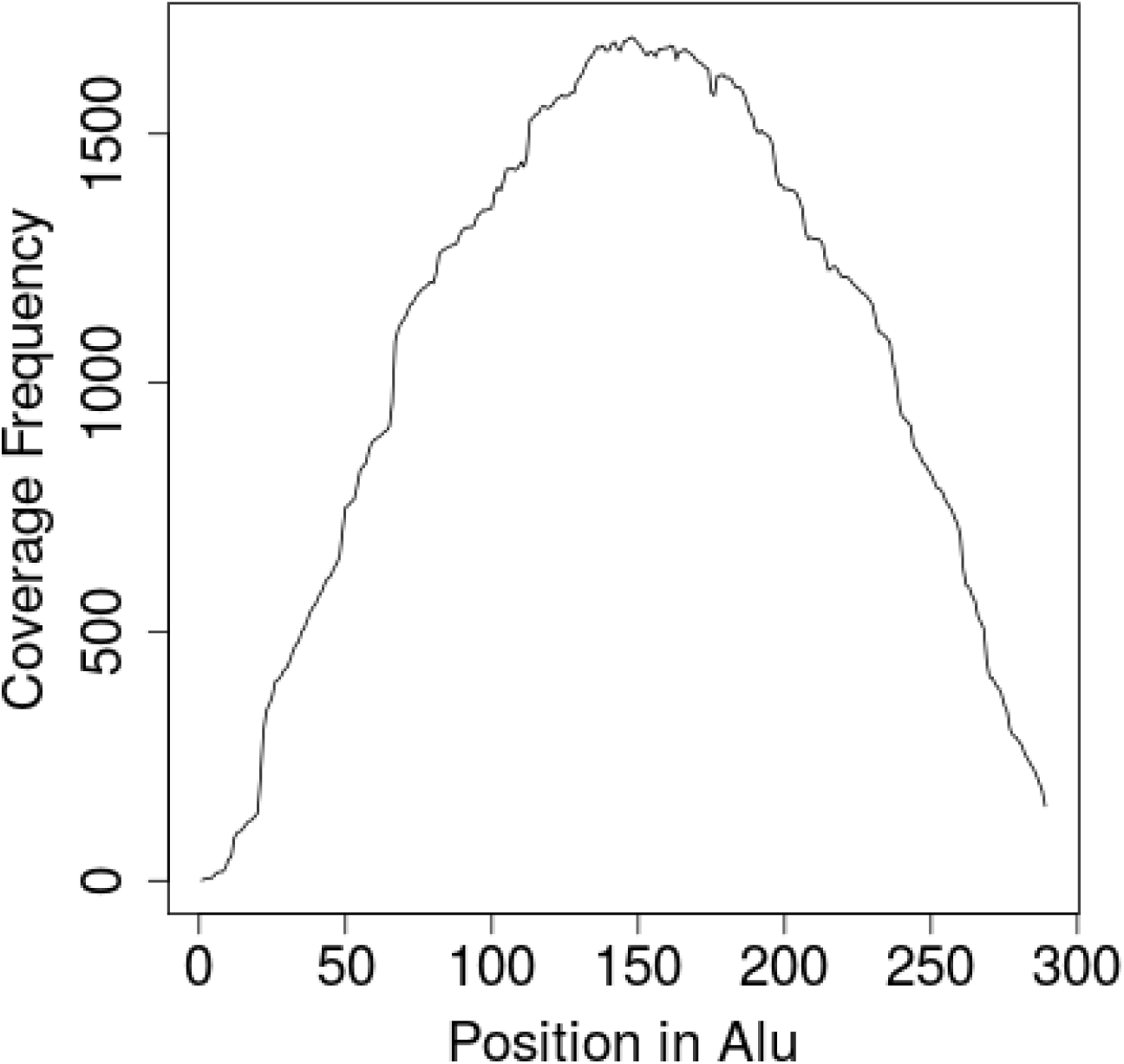
Coverage of the *Alu* Sequence by Conversion Events. For each site in *Alu*, the number of times it was included in a conversion tract was plotted.

Detecting conversion events depends on the number of differences between donor and recipient elements; conversion between more distantly-related elements is easier to detect. Long conversion tracts from younger to older elements are also easier to detect than the reverse, because older donors with longer tracts are easier to distinguish. The above analysis suggests, however, that the false negative rate is low (<5%) for conversion tracts larger than 100 bp, and so to evaluate possible sampling bias, we constructed a subset of 1,428 events with tracts 100 bp or larger. Elements from the youngest major *Alu* subfamily, *Alu*Y, are disproportionately likely to be both gene conversion recipients and donors in either the full conversion set or the large-tract subset (Table 2). The middle family, *Alu*S, is also disproportionately present as both recipients and donors, though to a lesser extent. Elements from *Alu*J, the oldest major subfamily, are greatly underrepresented as donors and recipients relative to their frequency in the genome. Thus, older elements are less likely than younger elements to be involved in gene conversion as either donors or recipients. This pattern runs opposite to the expected bias in identifiability, because older elements are more dissimilar to their potential conversion partners, but is expected from the strong relationship between sequence similarity and conversion probability^13^. Consistent with this interpretation, identified conversion pairs are more likely to belong to the same family than expected from the subfamily donor and recipient frequencies alone (Table 3).

**Table 2.**

|  | Recipients | Donors | Large-Tract Recipients | Large-Tract Donors | All gorilla Elements | Conversion Probability |
| --- | --- | --- | --- | --- | --- | --- |
| *AluY* | 18.0% (462) | 13.0% (334) | 21.6% (315) | 14% (205) | 8.6% (21,813) | 2.1% |
| *AluS* | 77.9% (2002) | 78.3% (2014) | 75.2% (1095) | 76% (1109) | 67.3% (171,041) | 1.2% |
| *AluJ* | 4.1% (105) | 8.5% (219) | 3.2% (46) | 9.5% (138) | 24% (61,010) | 0.2% |

**Table 3.**

| Recipient: Donor | Expected | Observed |
| --- | --- | --- |
| Y:Y | 2.3% | 3.7% |
| Y:S | 14.1% | 13.2% |
| Y:J | 1.5% | 1.1% |
| S:Y | 10.1% | 9.1% |
| S:S | 61.0% | 62.3% |
| S:J | 6.6% | 6.4% |
| J:Y | 0.5% | 0.2% |
| J:S | 3.2% | 2.8% |
| J:J | 0.3% | 1.0% |

Younger subfamily recipients are overrepresented more strongly than donors (Fisher exact test, p<0.0001) in both the overall dataset and the large-tract subset. This may be driven by a higher frequency of intact putative PRDM9 binding motifs, which strongly affect the probability of being a recipient but not a donor. Only 28% of *Alu*J elements have a fully intact motif, compared to 34% of *Alu*S elements and 68% of *Alu*Y elements. However, it is likely that sites on the *Alu* sequence other than the motif itself are important for double-strand break targeting^8^; supporting this, there is still a significant bias towards *Alu*Y in recipients versus donors even if both elements have fully intact motifs (Fisher exact test, p=.0013).

On average, pairs of converted loci differed at 53.9 sites (standard deviation 11.5 sites) in their ancestral state at the Great Ape root (the last point where we can infer ancestral sequences). In contrast, *Alu* elements overall differed by an average 67.3 positions. However, part of this is due to the bias towards younger elements in conversion pairs, and conversion pairs were only slightly more similar to each other than pairs of elements chosen at random from the recipient and donor pools (average difference 55.2 sites). Though highly significant in both the long-tract and full datasets (p=0.001, t-test), a difference of only 1.3 base pairs suggests that sequence similarity is surprisingly unimportant, at least for the level of interlocus divergence seen in this dataset. The sequence similarity between conversion pairs in our dataset is considerably lower than reported for other identified conversion events in primate genomes, which are generally above 95%^14^. The high density of *Alu* elements probably offered many more opportunities for pairwise interaction than the typical gene family, and it appears that opportunity based on distance along the genome was a far more important factor.

Conversion and post-conversion substitution events can be inferred by comparison of donor and recipient sequences with orthologous sequences at the root of the Great Apes. The average conversion event involved 21.4 identifiable conversion tract substitutions from the donor to the recipient element, compared to 3.7 substitutions along the entire terminal gorilla lineage at non-converting loci, implying that identified conversion events were responsible for at least 9.4% of *Alu* substitutions along the gorilla lineage. Although both allelic and interlocus gene conversion bias towards GC variants has been observed in a wide variety of species^15,16,17^, we do not observe such a bias (Table 4).

**Table 4.**

| Substitution Type | Full Set Count | Full Set Reverse Count | Long-tract Subset Count | Long-Tract Subset Reverse Count |
| --- | --- | --- | --- | --- |
| C:G→A:T | 3409 | 3930 | 2554 | 2899 |
| C:G→T:A | 20180 | 17205 | 14736 | 12494 |
| C:G→G:C | 4152 |  | 3086 |  |
| A:T→T:A | 3842 |  | 2859 |  |
| CpG:GpC→TpA:ApT | 6561 | 3752 | 4897 | 2745 |
| C:G→T:A (non-CpG) | 13619 | 13453 | 9839 | 9749 |

A relationship between TEs and PRDM9 binding motifs was previously noted for modern humans, in which THE1 elements appear associated with recombination hotspots^8^. Because our results suggest a PRDM9-binding site in gorilla *Alu*s, it seemed worthwhile to consider the relationship between PRDM9 and repetitive elements more generally. To do this we identified additional putative PRDM9 binding sites that appeared to be active in the human lineage by making use of the tendency for PRDM9 binding motifs to become depleted from the genome over time^18^. We scanned the genome for 12-mers that were depleted in humans relative to other terminal branches and identified 1,546 depleted 12-mers (p<10^-16^). Grouping similar or adjacent 12-mers produced 19 motifs (Supplementary Table 1). To verify whether the depleted motifs were PRDM9-binding sites, we looked for a signature of recombination based on an elevated rate of A➔C and A➔G substitutions that would result from GC-biased allelic gene conversion.^3^ The 100 bp flanking regions of depleted motifs show substantially elevated rates of A➔C and A➔G substitutions relative to the average genomic rate in human, but not of other substitution types or in other lineages, exactly as expected if these were PRDM9-binding motifs exclusively in the human lineage (Supplementary Figure 3). Interestingly, a number of the 19 groups can be speculatively aligned into plausible higher order groups (Supplementary Table 1), which is compatible with the PRDM9 protein having evolved to have distinct but similar binding motifs over time. Five of the motifs are found in a specific class of repetitive elements in at least 25% of cases, including *hAT-Charlie, ERVL-MaLR*, and *Alu* (Supplementary Table 1). For *hAT-Charlie* and *ERVL-MaLR*, the motifs associated with that repeat are 2-3.8 times more likely to become depleted within the repeat than elsewhere in the genome, suggesting that PRDM9 bound the motif more strongly in the background of the repeat. Based on this, we conclude that PRDM9 often targets motifs in repetitive sequences.

Overall, our observations are highly consistent with the idea that a burst of *Alu-Alu* recombination along the gorilla lineage was driven by PRDM9 recognition of a target site common among *Alu* sequences, resulting in an epidemic of sequence deletion in the gorilla genome. This raises the question why PRDM9 appears to target repeats, given the considerable risks associated with interlocus recombination, as exemplified in many human diseases.^6,19,20^. *Alu*, as a relatively young TE family with over a million copies in the gorilla genome, would appear to be a particularly dangerous target.^21^ Perhaps, as suggested previously^22^, the risks of interlocus recombination from targeting repetitive sequence are balanced by the benefits of avoiding targeting functional regions for double-strand breaks and associated mutations. In any case, our results suggest that genome instability driven by transient PRDM9 interaction with repeats is a recurring threat for mammalian genomes.

## Methods

### Obtaining *Alu* Ortholog Alignments

A dataset of *Alu* elements was obtained from the RepeatMasker^23^ annotation of the hg38 assembly of the human genome. Only full-length sequences (275-325 bp) were included, producing a dataset of 779,310 elements. Human elements were aligned to a consensus of the *Alu* sequence using the probabilistic version of the Needleman-Wunch algorithm described by Zhu et al.^24^

The 6-primate EPO^25^ whole-genome alignment was acquired from Ensembl release 71^26^, which is based on the GRCh37 assembly of the human genome. We used five of these genomes: human, chimpanzee, gorilla, orangutan and macaque. The positions of elements in the *Alu* dataset were used to identify *Alu* positions in the whole-genome alignment. Because EPO contains an alignment of orthologs to the human genome, we used these alignments, together with our alignments of human *Alu* to the consensus, to obtain alignments of each ortholog to the consensus.

We filtered out all *Alu* loci that did not include human, chimpanzee, gorilla, and orangutan orthologs. As we use macaque only for substitution rate estimation, we did not exclude loci missing only macaque. To protect against misalignment of *Alu*, we also filtered out loci in which chimpanzee, gorilla, or orangutan in either 100 bp flanking region around the element differed by more than 10% from the human flanking region.

### Testing Potential Conversion Pairs

Each pair of elements was tested for conversion between each element within 100Kb, considering each in turn as potential donor and recipient, and considering separately the possibility of conversion on each branch in the Great Ape phylogeny (i.e., the phylogeny including human, chimpanzee, gorilla, and orangutan as leaves).

Given a possible donor element, a possible recipient element, and a possible branch on which a conversion event occurred, we consider the relative likelihood between two scenarios: first, that the two elements evolved independently, neither experiencing gene conversion; second, that there was a conversion event in which a segment of the recipient element was completely converted to that of the donor element.

The marginal likelihood of a conversion event involving a specified potential recipient, donor, and branch is the sum of the likelihoods of each possible conversion tract. We thus estimate the likelihood of every possible contiguous tract in the alignment, relative to the scenario of no conversion. For each position in the tract, we consider two possible pairs of trees, the first indicating that the two homologous nucleotides at the possible donor and recipient evolved independently according to the normal Great Ape phylogeny, and the second indicating that the recipient branched off the donor at the point of conversion (Supplementary Figure 4). The relative likelihood of a tract position is the relative likelihood between the trees indicating non-independent evolution of the homologous nucleotides at that position and the trees indicating independent evolution. Outside of the tract, positions evolve along the same trees in the conversion and non-conversion scenario, so the relative likelihood of each position is one. The relatively likelihood of a tract is the product of the relative likelihood of each individual position in the tract. Using this approach, estimating the conversion likelihood reduces to the problem of estimating the relative likelihood of two sets of trees for every site in the tract.

Given a tree, substitution probabilities across each branch, and a set of leaves, the tree likelihood can be calculated using Felsenstein’s algorithm.^27^ The leaves of each tree were obtained from the homologous site in the element across the potential donor and recipient element in human, chimpanzee, gorilla, and orangutan. Substitution probabilities across the branches of the Great Ape phylogeny were straightforward to obtain, as described below. Estimating substitution probabilities on the branches surrounding the conversion event itself is more complicated, as the time of conversion is unknown. To address the unknown conversion timing, we averaged tract likelihoods across 100 possible conversion points spaced equally across the branch of the Great Ape phylogeny under consideration.

The conversion likelihood for a recipient-donor pair reflects the strength of evidence for conversion of recipient by donor relative to the possibility of independent evolution of each element without conversion. However, these are not the only possible scenarios. In particular, if the element under consideration as recipient was converted, but by a different donor, the conversion scenario may be strongly favored over the independence scenario for many considered donors that are closer to the actual donor than the recipient. To guard against such false positives, for every potential conversion event passing a likelihood threshold, we compared the donor-recipient pair under consideration to 10,000 pairings of the recipient with random elements in our dataset. Then, we determined the proportion of donors with higher likelihoods than the donor under consideration. Every donor-recipient pair, then, was associated with two values: the relative likelihood and donor percentile. We required, for a potential conversion pair to go into the conversion set, that the pair have relative likelihoods larger than e^30^, and the potential donor have higher likelihoods with the recipient than 99.95% of random potential donors. We chose these values to obtain a false positive rate below 5%, as described below.

If a donor-recipient pair was in the conversion set, it was of interest to infer properties of the conversion event, as well as information about the donor and recipient before and after the event. We drew events (given donor, recipient, and branch) from the posterior distribution using the following procedure. We first drew a tract with probability proportional to its relative likelihood. We then drew a point on the branch at which the event occurred, conditional on the selected tract. Conditional on the above, we then drew trees from the posterior using Felsenstein’s algorithm, with each tree indicating the nucleotide at each node in each species in the donor and recipient. For analyses, we drew one event from the posterior of each pair in the conversion set.

### False Positive Rate Estimation

To estimate false positive rates, for every element in the *Alu* alignment dataset dataset, we randomly placed it somewhere else in the genome, at least 1 Mb away from its true position. Then, we tested it as a potential conversion recipient for all elements within 100 Kb. We identify all events passing the same thresholds for admission to the conversion set, putting them in the false-positive set. Because large-distance conversion events are unlikely, these events should consist primarily of false positives. We estimated the false positive rate as the number of events in the false positive set divided by the number of events in the conversion set.

### Substitution Probability Estimation

Substitution probabilities were estimated from the EPO alignment, which contains inferred ancestral sequences. For each branch of the Great Ape phylogeny, the probability of being in state Y given state X at the start of the branch was estimated by dividing the number of X to Y changes by the number of positions in state X across all elements at the start of the branch. Substitution probabilities for each possible X to Y substitution were estimated separately, and we distinguished C to A and G to T substitutions at CpG sites from other such substitutions.

For most purposes, we could use these substitution probabilities across the Great Ape phylogeny directly. Given a conversion event along a branch, however, substitution probabilities needed to be estimated up to that event, and from that event to the present. To do this, we first converted our substitution probabilities to rates. As substitution probabilities are low, we assumed no more than one event per site, in which case:

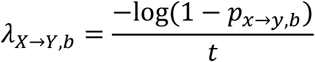
where *A_x→y,b_* is the rate of substitution from X to Y along branch b, *p_x→y_* is the probability of substituting from X to Y across that branch, and t is the branch length, defined to be 1 for the entire branch. After estimating the rates across the entire branch, we used this equation to estimate substitution probabilities along sections of the branch.

### Identifying *Alu* Recombination-Mediated Deletions

If an ARMD event occurs on a given branch of the Great Ape phylogeny, we expect to observe a single element in the genome in which that event occurred that is a hybrid between the two recombination participants. The event can be identified and characterized by comparing this hybrid element to the orthologs of the recombination participants. The upstream flanking region of a recombination product will match one participant and the downstream flanking region will match the other.^6^ We took locations of all *Alu* elements in the human, chimpanzee, gorilla and orangutan genomes from the RepeatMasker^23^ database and extracted the 100 bp flanking regions of each element in the hg38 assembly of the human genome, the panTro4 assembly of chimpanzee^28^, the gorGor3 assembly of gorilla^29^, and the ponAbe2 assembly of orangutan^30^. For each element in each genome, we attempted to match the upstream and downstream flanking regions to flanking regions of elements in the other genomes. To declare a match, we required at least 90% sequence identity using the EMBOSS^31^ implementation of the Smith-Waterman algorithm.

For each pair of genomes, we identified elements in which the upstream and downstream flanking sequences matched the upstream and downstream sequences of distinct elements in the other genome. To restrict our analysis to straightforward cases, we required that the upstream and downstream matches were on the same chromosomes and within 100kb of each other. For apparent recombinant elements that had both upstream and downstream matches in all three other genomes, we used these matches to perform probabilistic ancestral reconstruction of both recombination participants at the root of the Great Apes, using the same method as described above for gene conversion. For analysis, we drew a single reconstructed ancestral sequence from the posterior for each element involved in a recombination event.

### Identifying Ancient PRDM9 Motifs Using Depletion

To identify putative PRDM9 motifs, we examined 12-mers that were aligned in the 6-primate EPO alignment^26^ for each terminal branch of the Great Ape phylogeny (human, chimpanzee, gorilla, orangutan). For each of the 4^12^ 12-mers, we counted the totals that were either conserved among the four Great Apes, or that were conserved among three Great Apes and different in the fourth. Focusing on the detection of excess rates of depletion in the human lineage relative to the others, we considered only 12-mers that had depletion counts in human at least twice that of chimpanzee, and had at least 200 copies in the chimpanzee genome.

To detect significant depletion, we assumed that the number of depletions in chimpanzee, gorilla, and orangutan were Poisson distributed according to a constant per-motif rate λ_i_ for each motif *i*, weighted by the branch length, which was estimated as the average depletion count for each lineage across motifs. We then used the depletion counts at each motif *i* to calculated the probability distribution of λ_i_ under a uniform prior. We then calculated the probability of observing the number of depletions observed or greater in the human lineage under the assumption that it was also distributed according to λ_i_, weighted by the human branch length. We determined an element was depleted if this probability was less than 10^-16^. Note that the total number of 12-mers, allowing for ambiguous bases, is only 5^12^, so we do not expect to observe false positives as long as our assumptions are met. We first identified 12-mers that were significantly depleted individually, and subsequently examined merged counts for 12-mers including each single ambiguous base *N* to obtain greater power to detect significance.

After significance testing, we grouped together 12-mers that were aligned but differed slightly. We used an iterative strategy, starting with the most frequent 12-mer as a potential group founder and continuing to the next most frequent 12-mer that had not yet been grouped. For each potential founder, we considered each ungrouped 12-mer as a possible addition. We allowed no more than one mismatch of nucleotides (not including Ns), and no more than four of either mismatches or Ns in the founder or potential addition. In the next step, we joined groups if the founding members of each cluster had perfect overlap at 11 bases but differed at the ends, thus extending the possible length of motifs.

After grouping 12-mers, we searched the genome for each instance (depleted or not) of each significantly depleted 12-mer, collecting the 100 bp flanking regions. Substitution probabilities in flanking regions were estimated as described above for *Alu* elements, as were genome-averaged substitution probabilities. The RepeatMasker^23^ annotation of each 12-mer in the genome was also identified using the RepeatMasker Open-4.0.5 repeat libraries.

## Acknowledgements

We thank Andrew Martin and Kenji Fukushima for helpful discussions. We acknowledge the support of the National Institutes of Health (NIH; GM083127 and GM097251) to DDP.

---

**Supplementary Figure 1.**
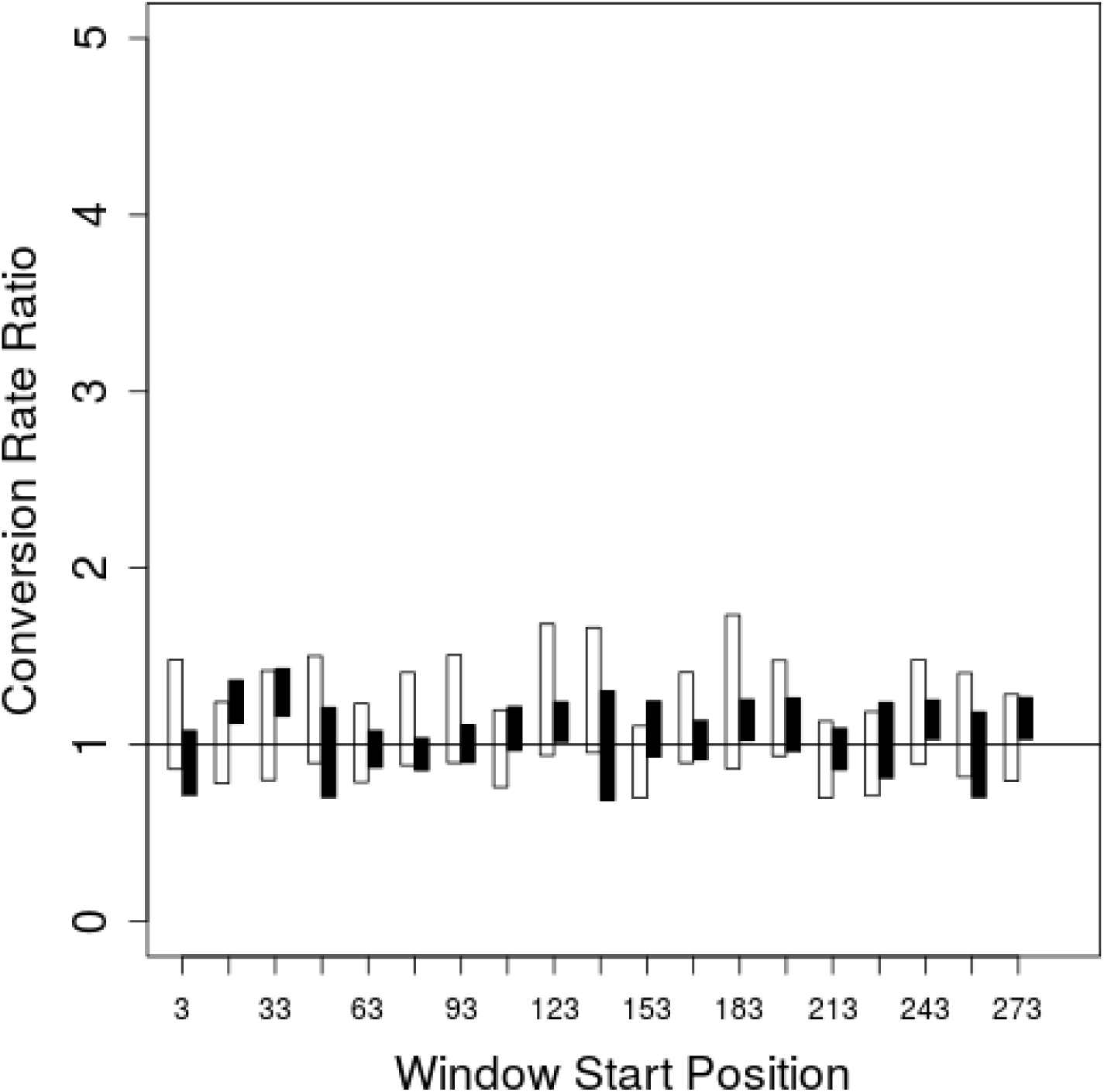
Conversion Donor Ratio Between Elements With and Without Perfect Match to the Consensus in 15 bp Windows Across *Alu*. For 15 bp windows across *Alu*, the proportion of elements that were donors in the gorilla gene conversion set was calculated for loci that had perfect match to the most common sequence at the root of the Great Apes in that window (excluding positions differing between *Alu*S, *Alu*Y, and *Alu*J) and all other elements. The white bars show 95% credible regions for AluY elements, while the black bars show 95% credible regions for *Alu*S elements.

**Supplementary Figure 2.**
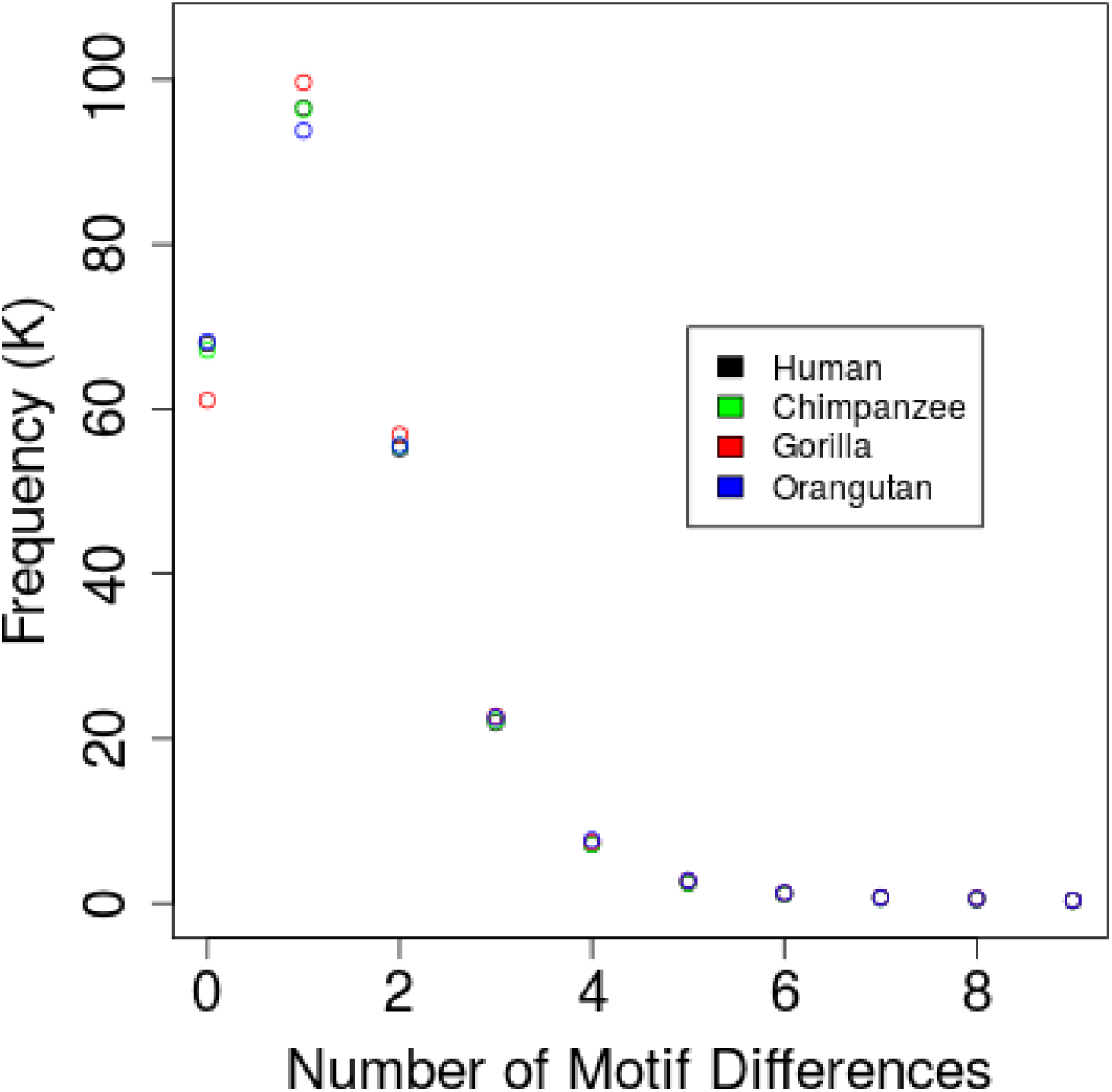
Differences from the Putative PRDM9 Motif among *Alu* Elements in the Great Apes. The frequency of elements with each count of differences from the putative PRDM9 motif is plotted for human, chimpanzee, gorilla, and orangutan.

**Supplementary Figure 3.**
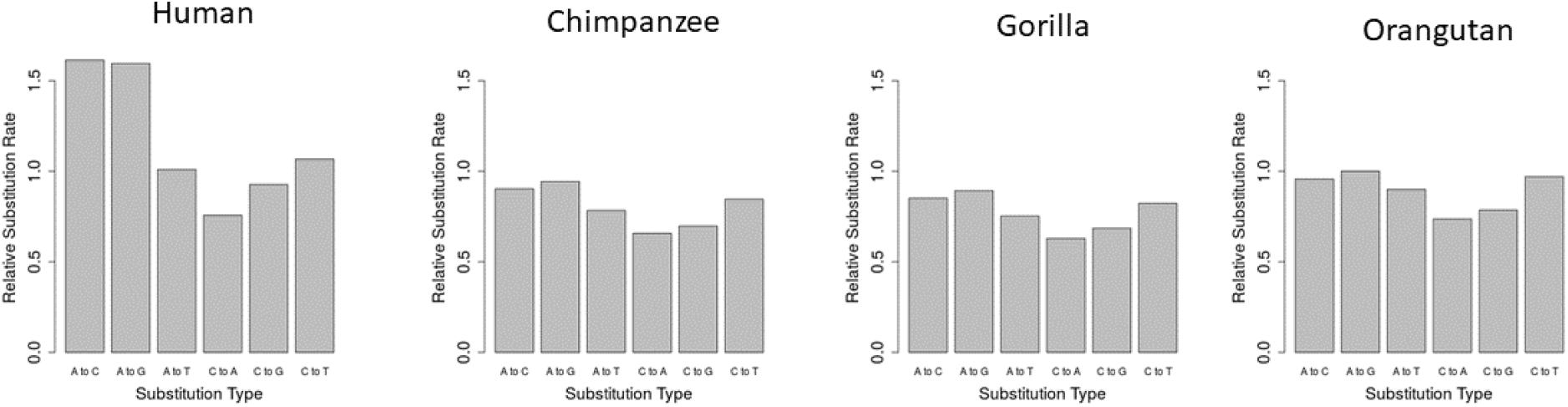
Substitution Rates Relative to Genome Average around Motifs Depleted in the Human Lineage. The substitution rate in each lineage was estimated in the 100 bp flanking regions of 12-mers depleted in the human lineage. These rates are plotted relative to the genome average, estimated from the entire whole-genome alignment.

**Supplementary Figure 4.**
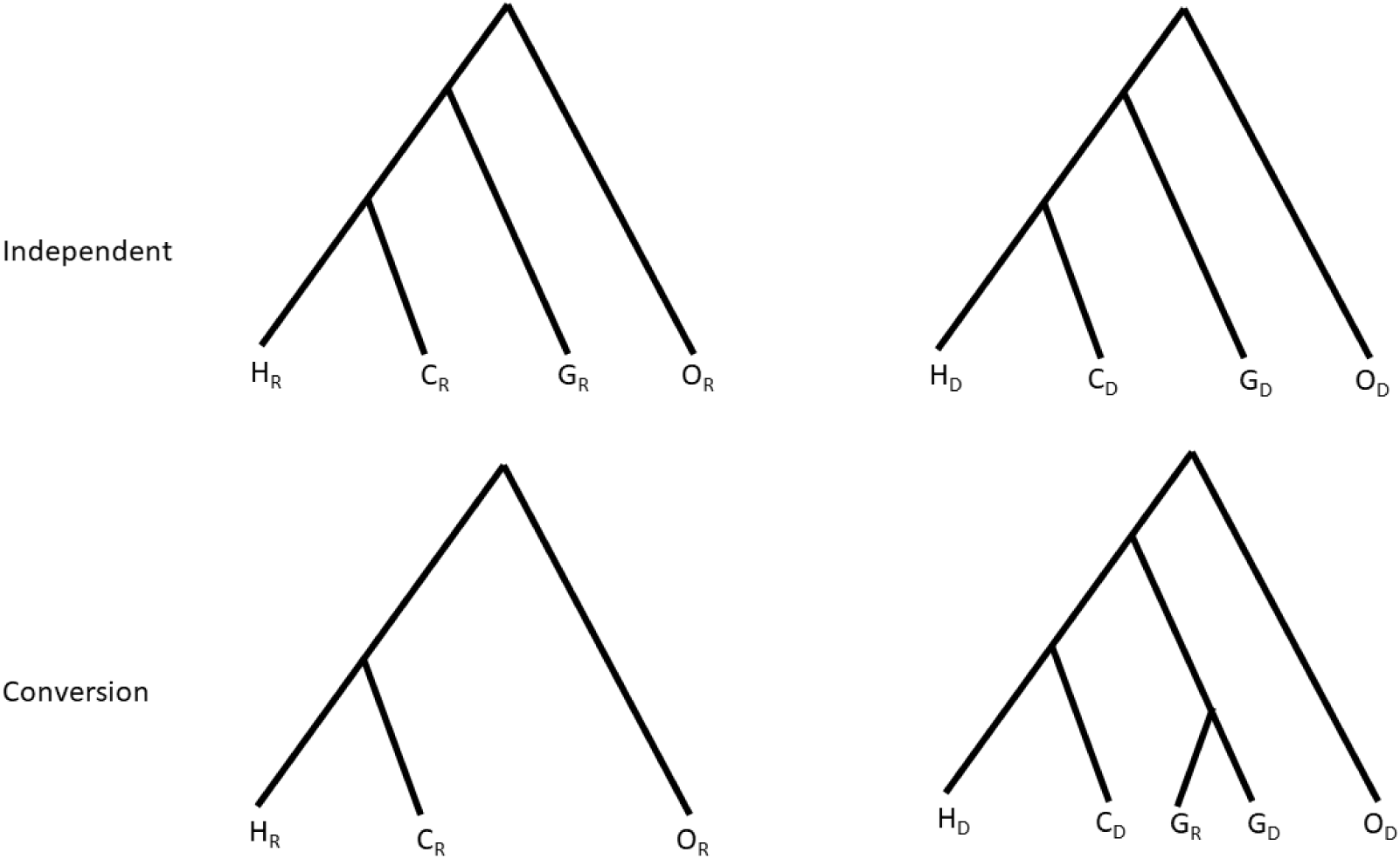
Trees representing scenarios of conversion and independent evolution for a homologous site at a potential recipient and donor locus. Each leaf represents one of eight homologous nucleotides among four orthologues and two loci. Leaf labels indicate species (human (H), chimpanzee (C), gorilla (G), and orangutan (O)) and presence at either the donor (D) or recipient (R) locus. Each set of trees represents one possible scenario. If the site was not converted, both sites evolved independently according to the typical Great Ape phylogeny. If the site was converted, the recipient branches off of the donor lineage after the conversion event. The tree shown corresponds to conversion in the gorilla lineage.

